# ZFHX4 is necessary for dopaminergic neuron differentiation and controls cell cycle by regulating LIN28A

**DOI:** 10.1101/2025.07.03.662987

**Authors:** Elena Valceschini, Borja Gomez Ramos, Jochen Ohnmacht, Aurelien Ginolhac, Marie Catillon, Deborah Gerard, Anthoula Gaigneaux, Dimitrios Kyriakis, Kamil Grzyb, Enrico Glaab, Anne Grünewald, Alexander Skupin, Thomas Sauter, Rejko Krüger, Lasse Sinkkonen

## Abstract

The selective degeneration of midbrain dopaminergic neurons (mDANs) is the main pathological hallmark of Parkinson’s disease (PD). Although many transcription factors (TFs) guiding mDAN development have been identified, the details of the underlying regulatory networks remain elusive. We have previously generated time-series transcriptomic and epigenomic profiles of human induced pluripotent stem cell (hiPSC)-derived mDANs. Integrative analysis of the data identified ZFHX4 as a prominent super-enhancer-controlled TF induced in mDAN differentiation. ZFHX4 has been associated with neurodevelopmental processes in several species and shows reduced expression in midbrain of PD patients. Using *in vitro* knockdown (KD) and overexpression experiments, we show that ZFHX4 is necessary but not sufficient for mDAN differentiation. ZFHX4 binds preferentially at active promoter regions and transcriptomic analysis upon ZFHX4 depletion during mDAN differentiation revealed putative primary target genes to be enriched for targets of cell-cycle-related TFs and pathways. Consistently, ZFHX4-depleted cells accumulated in G2-phase of the cell cycle, preventing normal cell cycle progression and exit. The RNA-binding protein LIN28A, involved in stem-cell maintenance and microRNA (miRNA) maturation, emerged as one of the most upregulated genes upon ZFHX4-KD, in parallel with downregulation of neurogenic miRNA miR-9. Moreover, the LIN28A locus was enriched for ZFHX4 binding in CUT&Tag analysis. Taken together, our analysis indicates a pivotal role for ZFHX4 in regulating the cell cycle, specifically in silencing multipotency and proliferative programs, while maintaining mDANs in a post-mitotic state by controlling LIN28A-miR-9 axis.

## Introduction

The midbrain dopaminergic neurons (mDANs) are anatomically divided into three regions, namely *Substantia Nigra pars compacta* (SNc), ventral tegmental area (VTA) and retrorubral field (RrF). mDANs located in the SNc mainly project to the striatum and degenerate in Parkinson’s disease (PD) ^1,2^. Notably, mDANs from the ventral tier of the SNc are known to be more susceptible to degeneration in PD compared to the ones from the dorsal tier ^3^. In contrast, mDANs from the VTA and RrF project to the ventral striatum and prefrontal cortex, respectively, and are involved in the control of motivation and reward ^4^. The neurons from these areas are less affected in PD but are associated to psychiatric disorders such as schizophrenia and drug addiction ^5,6^. Understanding the molecular mechanisms behind this selective vulnerability is therefore crucial for identifying the causes of the disease.

Transcription factors (TFs) together with signaling molecules play a pivotal role in cellular development and specification, defining context-specific regulatory programs. TFs bind to *cis*-regulatory elements of the genome, known as enhancers, and selectively recruit co-factors and co-activators to regulate gene expression ^7^. Enhancers regulating key aspects of cell identity are often clustered into dense regions called super-enhancer (SE), which are bound by multiple master regulators, leading to recruitment of high levels of the Mediator complex ^8^. These regions are particularly enriched in histone modifications such as histone H3 lysine 27 acetylation (H3K27ac), associated with active enhancers, and are characterized by exceptionally high levels of gene expression, and frequently harboring disease-associated genetic variants ^9,10^. SE-controlled TFs are thus central regulators of cell fate and cell identity, making their identification useful for understanding lineage specification processes.

Several TFs and morphogens controlling mDAN development have already been uncovered, including NR4A2 (also known as NURR1), one of the most prominent markers of mature mDANs, along with tyrosine hydroxylase (TH), the rate limiting enzyme for dopamine production ^11–14^. However, our current knowledge is primarily based on rodent models. To study human cells the use of induced pluripotent stem cell (iPSC)-derived models has become fundamental, given the low number and limited accessibility of human mDANs *in vivo* ^15,16^. Recent studies have demonstrated the benefits of iPSC-derived dopaminergic progenitor transplants in human PD patients, showing successful mDAN recovery, lasting up to several months ^17,18^. Furthermore, the advent of single-cell technology has paved the way for an unbiased identification of cellular subtypes based on their gene expression, allowing to identify novel subpopulations of mDANs, including those most affected in PD ^19,20^. Therefore, deploying these technologies will enable a better reconstruction of the molecular factors driving mDAN development and aid in uncovering novel cell-based therapies ^21^.

ZFHX4 is a transcription factor comprising 22 zinc fingers and four homeodomains whose expression is predominant in developing human brain and muscle ^22^, as well as in mouse cartilage and limb buds^23^. Pathogenic variants in ZFHX4 locus were shown to correlate with aberrant brain and craniofacial development, both in human and zebrafish models ^24,25^, while *Zfhx4*-deficient mice exhibit impaired endochondral ossification ^23^. Other than developmental processes, ZFHX4 has been shown to be detrimental for the survival of several types of cancers and the self-renewal of stem-cell like cells ^26–28^. In glioblastoma tumor initiating cells (TICs), for instance, its suppression led to an increase in neuronal markers, reduced cell proliferation and arrest in G0/G1 state ^29^. In TICs ZFHX4 was found to interact with CHD4, a core member of the nucleosome remodeling (NuRD) complex, involved in the regulation of cell cycle progression ^29^. While during palatal development, it partners with the TFs OSTERIX and RUNX2 ^23^. Furthermore, immunoprecipitation followed by mass spectrometry in neural precursor cells revealed that ZFHX4-interacting proteins are mainly involved in histone modifications, transcriptional regulation and development ^25^. Taken together, these data indicate ZFHX4 as a TF interacting with members of macromolecular complexes to exert context-specific regulation of both developmental and self-renewal pathways. Here we identified a novel role of ZFHX4 as a TF specifically induced during the *in vitro* differentiation of iPSC-derived mDANs. Depletion of ZFHX4 during neuronal differentiation led to a significant reduction in the number of mDANs while its overexpression did not lead to any changes in mDAN count. Identification of ZFHX4 target genes using CUT&Tag profiling and transcriptomic analysis upon ZFHX4 knock-down (KD) revealed its involvement in cell cycle regulation during mDAN differentiation, that was confirmed by immunocytochemistry and flow cytometry. Furthermore, LIN28A, a multipotency promoting factor controlling the maturation of the neurogenesis-promoting miR-9, appeared among the most affected primary targets of ZFHX4. In summary, ZFHX4 emerged as a TF necessary for mDAN development, regulating cell cycle by controlling the expression of targets like the miR-9-inhibiting multipotency factor LIN28A.

## Results

### ZFHX4 is a super-enhancer-controlled transcriptional regulator induced during dopaminergic neurogenesis

We have previously performed transcriptomic and epigenomic profiling across a differentiation time-series of iPSC-derived mDANs ^30^. Integrative analysis of the generated data enabled the reconstruction of gene regulatory networks at selected time-points and led to the identification of 49 SE-controlled TFs (Table S1)^30^. In parallel, we filtered all expressed TFs for those upregulated at each time-point of differentiation (log_2_-fold change >= 1, FDR < 0.05), resulting in a list of 33 TFs selectively expressed in mDANs (Table S2). These TFs were then compared with the 49 SE-controlled TFs, revealing 7 SE-controlled TFs with selective expression in mDANs (Supplementary Figure 1A). Among these putative key regulators of mDAN differentiation, ZFHX4 was found to be significantly downregulated in post-mortem samples of SNc from PD patients (Weighted average log_2_-fold change −0.51, FDR = 0.0021)^31^, making it a particular interesting regulator for further analysis.

LowC analysis of the *ZFHX4* locus in mDNAs revealed a >2 Mb topological domain with many regulatory interactions also from distal enhancers, carrying only one additional protein-coding gene (*PEX2*) and a few non-coding genes (Figure 1A) (Gerard *et al*, under revision). The progressive increase in accessibility at the ZFHX4 locus across the differentiation time-series - along with the broad enrichment of H3K27ac at its promoter - are characteristic of a SE-controlled target gene (Figure 1B). Analysis of epigenomic and transcriptomic data of 41 human tissues and cell types from EpiAtlas of International Human Epigenome Consortium (IHEC) revealed a high expression of ZFHX4 in neural cells and neural progenitors, indicating its involvement in the development and/or maintenance of neural cell identity (Figure 1C) ^32^. Consistently, highly transcriptionally active chromatin state was observed only in two samples, brain and hepatocytes. Single-nuclei transcriptomic analysis of post-mortem human SNc further shows a preferential expression of ZFHX4 in mDANs compared to other cell types in the same brain region (Figure 1D) ^19^. The selective expression of ZFHX4 and induction during mDAN differentiation were also confirmed in alternative *in vitro* models of iPSC-derived mDANs through time series analysis both at single cell (Figure 1E) and at bulk level (Figure 1F), and across tens of iPSC lines (Figure 1G) ^33,34^. Together, these data suggest that ZFHX4 is a SE-controlled TF, whose function remains largely unexplored and that seems to play a selective role in the context of mDAN differentiation.

**Figure 1:**
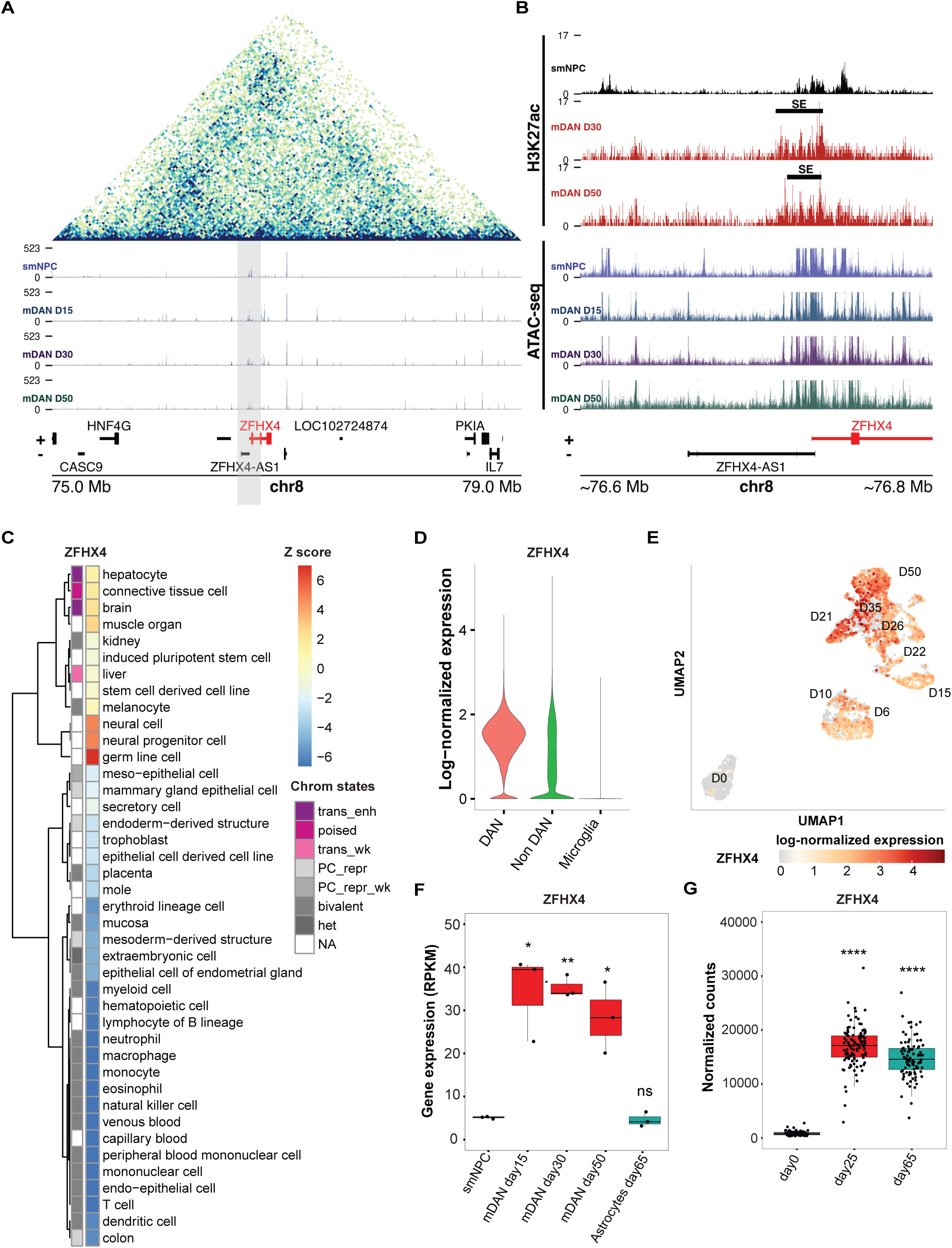
Transcriptomic and epigenomic profile of ZFHX4 in different models of mDANs. A) Topological associated domain at *ZFHX4* locus in 30 days differentiated mDANs and identified accessible chromatin regions across indicated time points. B) H3K27ac signal and chromatin accessibility at 200 kb region flanking *ZFHX4* TSS at indicated time-points of mDAN differentiation. Black bar indicates the identified mDAN-specific SE. C) Heatmap showing the expression and the ChromGene ^58^ state of ZFHX4 in 41 human tissues. D) Expression of ZFHX4 in selected cell-types DAN (TH^+^, SLC6A3^+^, SLC18A2^+^), Non-DAN (TH^-^, SLC6A3^-^, SLC18A2^-^) and microglia (TMEM119^+^, AIF1^+^, ITGAM^+^) from single-nuclei RNA-seq analysis of post-mortem *in vivo* human SNc ^19^. E) UMAP showing the log-normalized expression of ZFHX4 during time-series *in vitro* differentiation of iPSCs into mature day 50 mDANs, upon single-cell RNA-seq analysis. F) RNA-seq expression levels of ZFHX4 during time-series differentiation from smNPCs and in smNPC-derived astrocytes (N = 3). G) Expression of ZFHX4 in 95 different iPSC lines differentiated into mDANs ^33^. Error bars correspond to ±1 standard deviation (SD) from the mean, *t*-test, * = p-value < 0.05, ** = p-value < 0.01, *** = p-value < 0.001, **** = p-value < 0.0001, and ns = not significant.

### ZFHX4 is necessary but not sufficient for mDAN differentiation

To investigate the role of ZFHX4 in mDAN differentiation, we performed knockdown (KD) experiments at two selected time-points of differentiation (day 1 and day 9) to achieve both an early and a late KD (Figure 2A). Transduction with lentiviral vectors carrying shRNA targeting ZFHX4 resulted in an approximately 20% reduction of ZFHX4 expression in mDANs at day 15 compared to control shRNA (shSCRAMBLE) (Figure 2B). To study the impact of ZFHX4 depletion on mDAN numbers, neurons were analyzed using Fluorescence activated cell sorting (FACS) on day of analysis, taking advantage of the TH-mCherry reporter system that allows mCherry signal only in TH expressing cells ^30^. We observed a ∼60% reduction in mDANs following early KD of ZFHX4 and ∼40% following late KD, suggesting ZFHX4 is necessary for mDAN development (Figure 2C).

**Figure 2:**
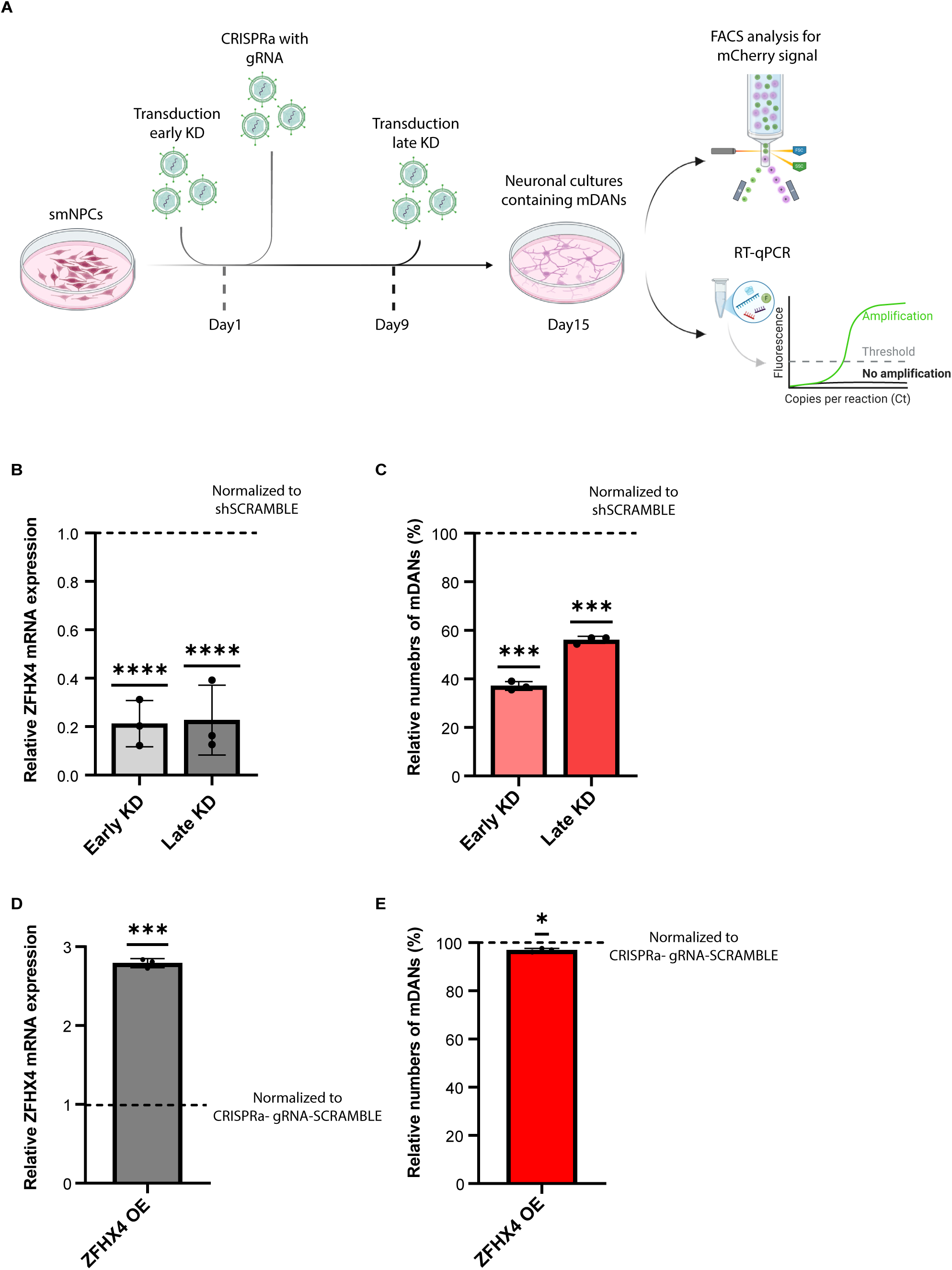
ZFHX4 is necessary but not sufficient for mDAN differentiation. A) Schematic representation of the early and late KD experiments and the CRISPRa for overexpression. FACS and RT-qPCR analyses were conducted on day 15. B) Early and Late transduction results showing ZFHX4 KD expression levels. C) Early and Late transduction results showing the effect of the KD on mDAN numbers. D) Results of ZFHX4 overexpression levels. E) Effect on mDAN numbers upon ZFHX4 overexpression. Data in all panels are representative of at least 3 independent experiments. Error bars (B-E) correspond to ±1 standard deviation (SD) from the mean. One sample t-test was used for statistical analysis, taking 100 (C, E) or 1 (B, D) as the theoretical mean. * = p-value < 0.05, ** = p-value < 0.01, *** = p-value < 0.001, **** = p-value < 0.0001, and ns = not significant.

To gain further insights into the role of ZFHX4 in mDAN differentiation, we overexpressed ZFHX4 following the experimental scheme shown in Figure 2A. Using CRISPR activation (CRISPRa) an almost three-fold overexpression of ZFHX4 (Figure 2D) was achieved. However, this perturbation did not result in any changes in proportion of mDANs (Figure 2E). Altogether these results indicate that ZFHX4 is necessary for mDAN development, but it is not sufficient on its own to drive their differentiation.

### ZFHX4 binds preferentially to active promoters and controls genes involved in cell cycle regulation

To map the putative targets of ZFHX4 in mDANs, we performed CUT&Tag analysis of ZFHX4 and H3K27ac in FACS-sorted mCherry-expressing mDANs (Figure 3A). ZFHX4 showed preferential enrichment at gene promoters and transcription start sites, similarly to H3K27ac. To compare the two datasets, we visualized the CUT&Tag signal for both ZFHX4 and H3K27ac, ranking TSSs according to the ZFHX4 signal for the top 20,000 gene loci (Figure 2B, Table S3). The ZFHX4 bound loci were largely acetylated at H3K27 although the signal intensity did not follow an identical ranking.

**Figure 3:**
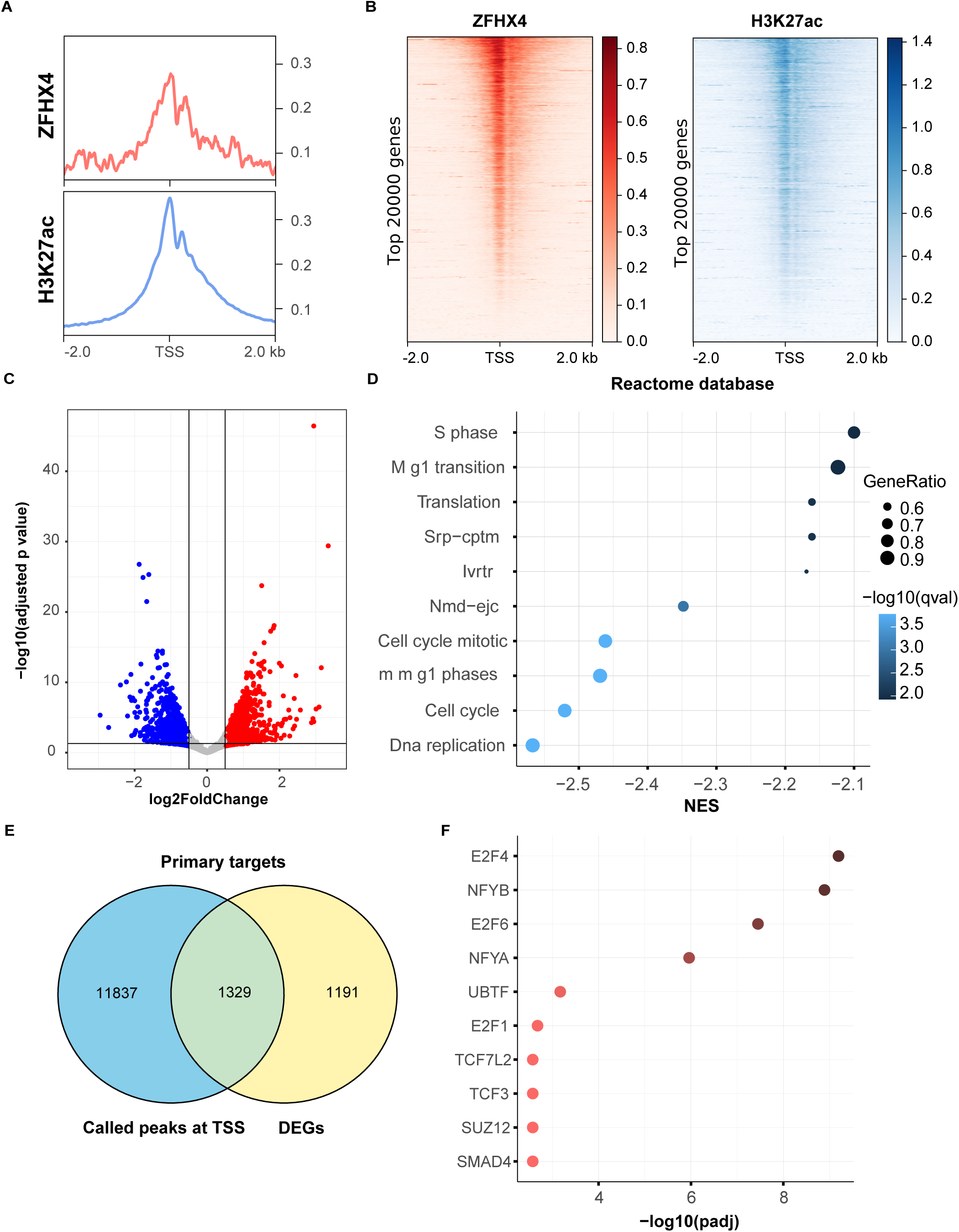
ZHFX4 is involved in the regulation of the cell-cycle. A) CUT&Tag profile of ZFHX4 and H3K27ac in day15 mDANs shows enriched signal at TSSs. B) Heatmap of ZFHX4 and H3K27ac CUT&Tag signal, ranked according to the top 20,000 ZFHX4-bound TSSs. C) Volcano plot of the 1,947 significant DEGs from the RNA-seq analysis of neurons on day 15 following a late transduction with shScramble or shZFHX4. Black lines represent cut-off according to FDR <0.05 and absolute log_2_-fold change >0.5. Red dots represent the upregulated genes and blue dots the downregulated genes. D) GSEA enrichment analysis of the 1,947 DEGs using the Reactome Database. X-axis represents the Normalized Enrichment Score (NES). E) Venn diagram of the intersection between the 13,116 called ZFHX4 peaks (log-q>19) from the CUT&Tag analysis and the 2,520 DEGs (FDR<0.05), leading to the identification of 1,329 putative primary targets of ZFHX4. F) Enrichment analysis using the ENCODE and ChEA Consensus TFs from ChIP-X database ^39,40^ on the 1,329 identified primary targets.

To further analyze the downstream targets of ZFHX4, we performed RNA-seq analysis following the late shRNA induced KD of ZFHX4 in mDANs, identifying 1,947 differentially expressed genes (DEGs) between shZFHX4 and shSCRAMBLE (absolute log_2_-fold change > 0.5, FDR < 0.05), with comparable number of up- and downregulated genes (Figure 3C, Table S4). Gene set enrichment analysis (GSEA) of the DEGs using the Reactome Database ^35,36^, revealed an enrichment of pathways related to the cell cycle, particularly S phase and transition from mitotic (M) phase to G1 phase (Figure 3D). To identify putative primary targets of ZFHX4, we intersected the list of DEGs with the CUT&Tag analysis-derived 13,166 ZFHX4-bound targets (log-q-value > 19) (Table S5). We then performed an enrichment analysis of these candidate primary targets using EnrichR on the ENCODE and ChEA Consensus TFs from ChIP-X database (Figure 3F)^37–40^. The results revealed a significant enrichment for the targets of E2F family TFs, key regulators of cell cycle progression, as well as of the NF-Y transcription factor complex, known for its roles in cell proliferation and developmental regulation ^41,42^. Together, these data suggest a role of ZFHX4 in the regulation of the cell-cycle dynamics during mDAN development.

### ZFHX4 depletion leads to cell cycle alterations in differentiating neuronal progenitors

Based on the observation that ZFHX4 appears to regulate cell-cycle genes during mDAN differentiation, we hypothesized that ZFHX4 might be involved in the cell-cycle exit during neurodevelopment, facilitating the transition from a proliferative state to a post-mitotic, quiescent state^43^. We therefore analyzed the proliferation state of the cells upon ZFHX4 KD.

First, we confirmed ZFHX4 depletion at protein level using immunocytochemistry, resulting in over 50% reduction on the day of analysis (Figure 4A-B). We then stained the cells for the proliferation marker Ki67, the mitosis marker phospho-histone H3 (PH3), and the neuronal marker MAP2. Upon ZFHX4 KD, we observed an increasing trend for both Ki67 and PH3, suggesting a rise in the number of proliferative cells when ZFHX4 was depleted (Figure 4A-D).

**Figure 4.**
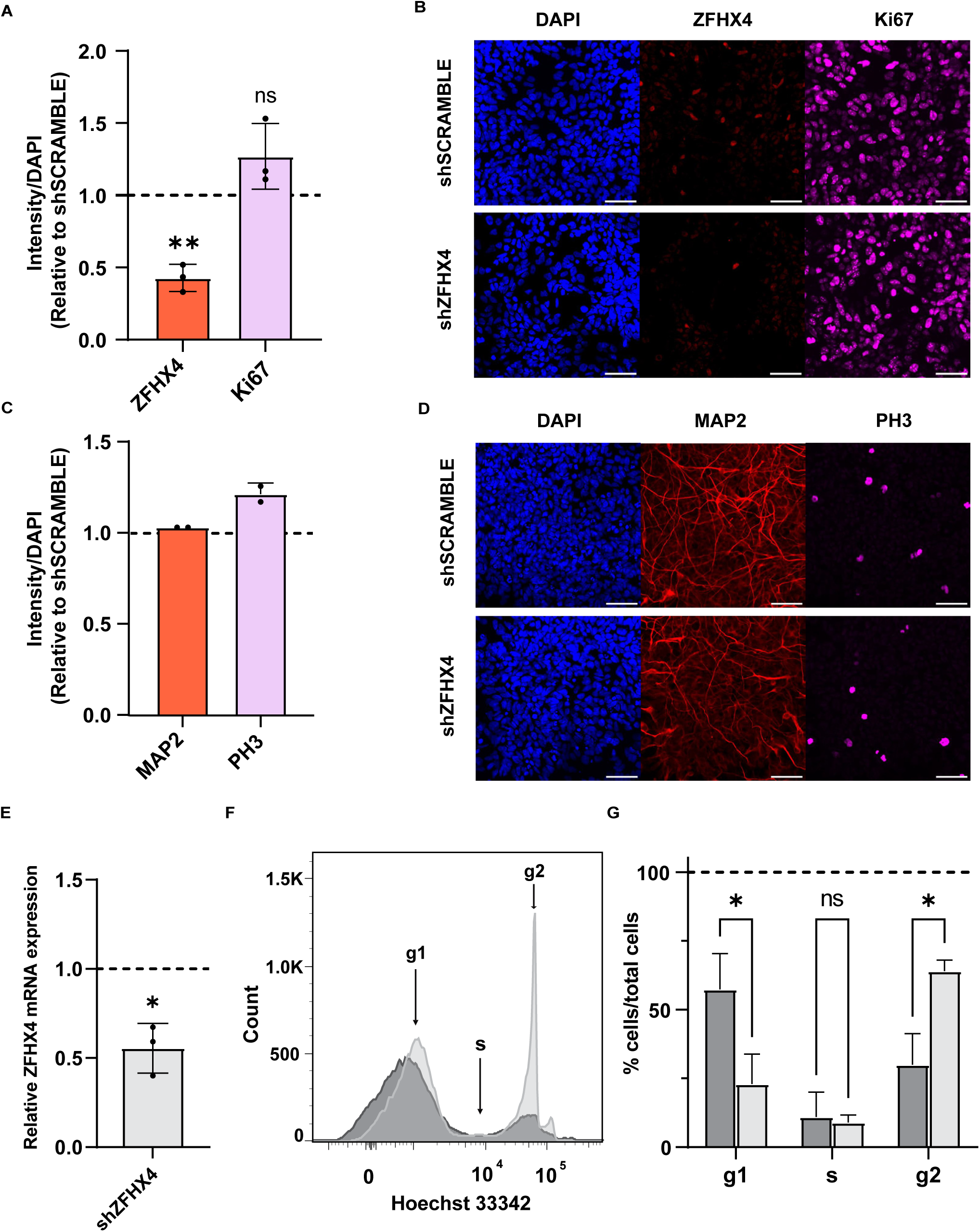
Failure to exit the cell-cycle is observed in the absence of ZFHX4. A) Quantification of the Ki67 or ZFHX4 stained area over the DAPI-stained area. Ratios were normalized to the shSCRAMBLE per replicate. Analysis was performed on day 8 following early ZFHX4 KD. (B) Representative images of the nuclear marker DAPI, ZFHX4 and the proliferation marker Ki67. C) Quantification of the MAP2 or PH3 stained area over the DAPI-stained area. Ratios were normalized to the shSCRAMBLE per replicate. Analysis was performed on day 8 following early ZFHX4 KD. (D) Representative images of the nuclear marker DAPI, the neuronal marker MAP2 and the mitosis marker PH3. Error bars (A, C) correspond to ±1 standard deviation (SD) from the mean, *t*-test, * = p-value < 0.05, ** = p-value < 0.01, *** = p-value < 0.001, **** = p-value < 0.0001, and ns = not significant. For Ki67 and ZFHX4 N = 3 while for MAP2 and PH3, N = 2. E) Expression level of ZFHX4 after early KD and RT-qPCR analysis on day 8. F) Hoechst 33342 positive cell counts after flow cytometry for either shZFHX4 (light grey) or shSCRAMBLE (dark grey). Analysis was done on day 8 upon early ZFHX4 KD. G) Quantification of cell counts according to each cell-cycle phase between shSCRAMBLE (light grey) and shZFHX4 (dark grey). Error bars correspond to ±1 standard deviation (SD) from the mean, *t*-test, * = p-value < 0.05, ** = p-value < 0.01, *** = p-value < 0.001, **** = p-value < 0.0001, and ns = not significant. (N = 3).

Next, we performed flow cytometry to further investigate the role of ZFHX4 in cell cycle-regulation. Early ZFHX4 KD was confirmed on the day of analysis by RT-qPCR, reaching approximately 50% reduction in expression. Staining with the DNA-binding fluorescent dye Hoechst 33342 revealed significant changes in the proportion of cells in the different cell-cycle phases (Figure 4F, Supplementary Figure 2). Specifically, there was an increase in the number of cells in G2-phase and a corresponding decrease in G1-phase cells, indicating a failure to exit and accumulation in the later stages of the cell cycle in the absence of ZFHX4 (Figure 4G). Altogether these data suggest that ZFHX4 is involved in regulating the cell cycle during mDAN development, possibly by facilitating the transition towards a mature state.

### ZFHX4 is necessary for repression of LIN28A and induction of neurogenic miR-9 during mDAN differentiation

To further characterize the downstream targets of ZFHX4, we analyzed the most significantly upregulated and downregulated genes following ZFHX4 KD. Among the most upregulated genes, we identified LIN28A (7.69-fold increase), while the primary microRNA (pri-miRNA) miR-9-3 was among the most downregulated genes (4.3-fold decrease) (Figure 5A). LIN28A is an RNA binding protein known to promote pluri- and multipotency, and has been previously shown to participate in an inhibitory loop with the neurogenesis-promoting miR-9 ^44^. We thus explored the transcriptomic and epigenomic profiles of these two factors in detail in our *in vitro* models (Figure 5B-C).

**Figure 5:**
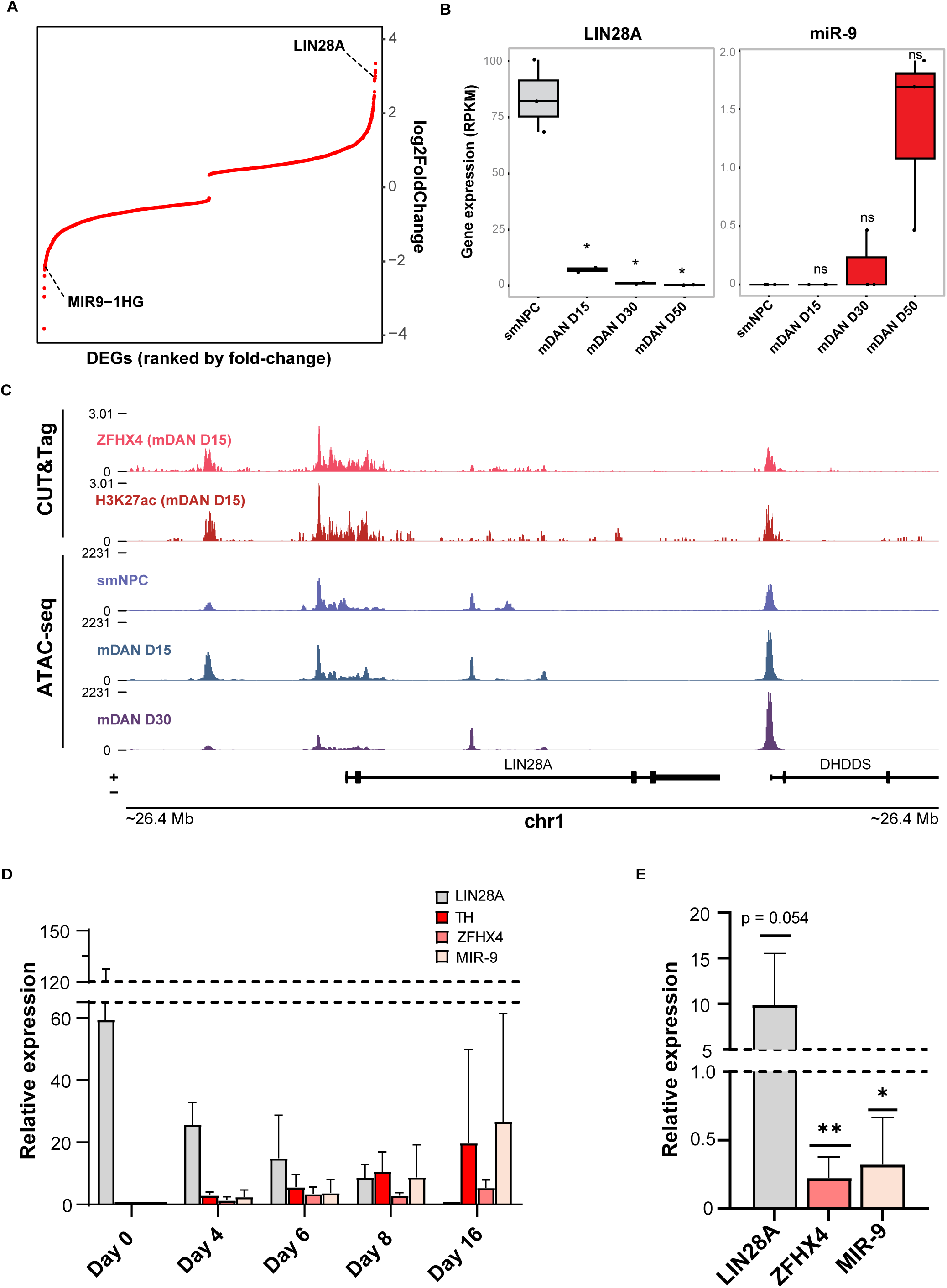
Differentiation failure is associated with ineffective repression of LIN28A. A) Rank plot of the 2,520 DEGs upon ZFHX4 KD highlighting LIN28A and miR-9 position. B) RNA-seq expression levels of LIN28A and pri-miR-9-3 during mDAN time-series differentiation (N = 3). C) CUT&Tag profile of ZFHX4 and H3K27ac at *LIN28A* locus. ATAC-seq profiles at selected time-points of *LIN28A* locus. D) Expression level of LIN28A, TH, ZFHX4, miR-9 after RT-qPCR and TaqMan during time-series mDAN dfifferentiation at selected time-points (N = 3). E) Expression levels of LIN28A, ZFHX4 and miR-9 upon late KD of ZFHX4 and analysis on day 15 (N = 3). Error bars correspond to ±1 standard deviation (SD) from the mean, *t*-test, * = p-value < 0.05, ** = p-value < 0.01, *** = p-value < 0.001, **** = p-value < 0.0001, and ns = not significant (B, E).

Time-series RNA-seq analysis confirmed that LIN28A is highly expressed in neural precursor cells (smNPC) and is progressively repressed during mDAN differentiation, whereas the neurogenic pri-miR-9-3 is induced during mDAN development. ATAC-seq analysis of the LIN28A locus showed a progressive decrease in chromatin accessibility, consistent with its transcriptional repression during mDAN differentiation (Figure 5C). Interestingly, CUT&Tag analysis revealed ZFHX4 enrichment at the LIN28A promoter, suggesting direct transcriptional regulation of LIN28A by ZFHX4 (Figure 5C). We then analyzed the expression of LIN28A, TH, ZFHX4 and mature miR-9 across multiple time points of early mDAN differentiation. The results confirmed the transcriptional repression of LIN28A during mDAN differentiation, concurrent with the induction of TH, ZFHX4 and miR-9 (Figure 5D). Finally, we validated the effects of ZFHX4 depletion using TaqMan microRNA assays and RT-qPCR, confirming an increase of LIN28A and the significant downregulation of mature miR-9 (Figure 5E). Altogether, these data indicate a role of ZFHX4 in the regulation of the LIN28A-miR-9 loop. Specifically, ZFHX4 appears to downregulate LIN28A, possibly directly at the transcriptional level, leading to its depletion during normal mDAN differentiation, in turn allowing the induction of neurogenesis-promoting miRNA miR-9, further supporting the suppression of cell cycle ^45,46^.

## Discussion

In this study, we identified ZFHX4 as a novel SE-controlled TF regulating mDAN differentiation. While ZFHX4 has previously been linked to neurodevelopmental processes, its role in mDAN development has not been described before. Mutations in the ZFHX4 locus have been associated with intellectual disability as well as craniofacial and brain developmental defects ^47^. ZFHX4 expression has also been observed in cartilage, where it has been demonstrated to play a role during endochondral ossification, suggesting its importance in developmental regulation ^23^. We have performed an integrative analysis of ATAC-seq, RNA-seq and Low input ChIP-seq data of *in vitro* iPSC-derived mDANs, identifying several SE-controlled TFs involved in mDAN differentiation ^30^. Among these, ZFHX4 emerged as a newly associated TF, with selective expression in mDANs and reduced expression *in vivo* in PD. Functional studies revealed that ZFHX4 depletion during mDAN differentiation leads to a significant reduction in proportion of mDANs, suggesting a necessary role during their development (Figure 1B-C). However, ZFHX4 overexpression was not sufficient to increase the number of mDANs (Figure 1D-E). The complex structure of ZFHX4 comprising 4 homeodomains and 22 zinc fingers suggests that it interacts with multiple co-factors to mediate context specific regulatory functions. Immunoprecipitation coupled with mass-spectrometry has identified ZFHX4-associated co-factors in different biological systems, including neural precursor cells and TICs ^29,47^. These findings support the notion that ZFHX4 requires multiple co-factors to exert context-specific roles. Therefore, overexpression on its own is probably not sufficient to increase the number of mDANs during differentiation.

Transcriptomic analysis following ZFHX4 KD revealed enrichment of cell-cycle related pathways in mDANs, particularly those involved in the S phase and the transition from mitotic (M) phase to G1 phase. Putative primary targets of ZFHX4, identified through CUT&Tag and RNA-seq analysis, were enriched for E2F target genes and components of the NF-Y transcription factor complex. The cell cycle is a tightly regulated process, with distinct checkpoints controlled by multiple factors to ensure proper cell division. E2Fs are a large family of TFs whose activity depends on the phosphorylation state of retinoblastoma proteins (pRB) and which sequentially regulate the expression of cell-cycle related genes during cell division ^48^. NF-Y is a heterotrimeric transcription factor complex that modulates the expression of genes involved in cell proliferation and developmental processes ^49^. During neurodevelopment, cells transition from a multipotent, proliferative state into a non-proliferative, quiescent state referred to as G0 phase, making their exit from the cell cycle division ^50^. The involvement of ZFHX4 in cell cycle regulation has previously been demonstrated in the context of glioma TICs where ZFHX4 suppression led to a decrease in proliferation and arrest in G0/G1 state ^29^. However, ZFHX4 deficiency in both mouse and zebrafish models resulted in a failure to differentiate into mature states ^23,25^. Moreover, in the context of neural differentiation, ZFHX4 has been shown to bind to transcription factors that promote cellular proliferation ^25^. Nevertheless, the regulation of the cell cycle during neurodevelopment, as well as the specific role of ZFHX4 during developmental processes, remains incompletely understood. We thus hypothesized that ZFHX4 may facilitate the transition from proliferative progenitor cells to mature neurons during mDAN differentiation. The analysis of the cell cycle state following ZFHX4 KD did not produce significant changes in the proliferation marker Ki67 nor in the mitotic marker PH3, although a trend toward increased Ki67 was observed (Figure 4A-D). Ki67 is a DNA-binding protein that displays distinct subnuclear localizations throughout all active phases of the cell-cycle and is absent only in the G0 phase. However it does not distinguish between G1, S G2/M phases ^51^. In contrast PH3 marks a post-translational modification of H3 that occurs during late G2 phase and mitosis, disappearing by anaphase, and therefore does not label cells in G1 or S phase ^52^. We thus decided to perform flow cytometry analysis using Hoechst 33342, a DNA fluorescent dye that enables discrimination of the G1, S, G2 phases based on the DNA content. The analysis revealed an increase in the number of cells in the G2 phase and corresponding decrease in the G1-phase cells following ZFHX4 KD (Figure 4E-G). These findings indicate that in the absence of ZFHX4, cells fail to exit the cell cycle and instead continue to proliferate or accumulate in late stages of cell cycle, thereby impairing proper development.

To further investigate the underlying mechanisms connecting ZFHX4 to mDAN differentiation and cell cycle control, we examined the putative primary targets of ZFXH4 based on our transcriptomic and epigenomic data. LIN28A was among the most upregulated genes upon ZFHX4 depletion (Figure 5A-B), and its promoter region was enriched for ZFHX4 signal in CUT&Tag, suggesting a direct transcriptional regulation by ZFHX4. The RNA-binding protein LIN28A is a known reprogramming factor implicated in early embryonic development and pluripotency ^53^. LIN28A participates in the post-transcriptional regulation of miRNA processing. Notably, during neurodevelopment LIN28A is known to inhibit the maturation of miR-9, a miRNA that in turn promotes neurogenesis. Consistently, miR-9 expression was significantly reduced (Figure 5B, E) upon ZFHX4 KD, supporting a regulatory axis involving ZFHX4, LIN28A, miR-9 ^44^. Hence, the strong upregulation of LIN28A and presence of ZFHX4 at LIN28A locus, together with its role in maintaining pluripotency, supports the hypothesis that ZFHX4 contributes to silencing of multipotency and proliferative programs, and promotes neurodevelopment.

In conclusion, we have identified a novel role for ZFHX4 as a SE-controlled TF essential for human mDAN development. ZFHX4 appears to be regulating the cell cycle by inhibiting multipotency programs, specifically by repressing the pluri- and multipotency-promoting factor LIN28A. This releases the expression of miR-9, promotes neurodevelopment and activates postmitotic programs.

## Materials and Methods

### Cell lines

The TH-Rep1 reporter cell line was derived from the human induced pluripotent stem cell (iPSC) line GM17602 (Coriell), which also served as a control during fluorescence-assisted cell sorting (FACS) in this study. This iPSC line has been previously utilized and described in previous studies ^30,54^, where it was used to establish a reporter system. To generate the TH-Rep1 line, a CRISPR/Cas9-based approach was employed to introduce a biallelic insertion of a T2A-mCherry sequence at the stop codon of the endogenous tyrosine hydroxylase (TH) gene. Routine testing confirmed that the cell lines remained free of mycoplasma contamination.

### Cell culture and differentiation

The protocol for differentiating small molecule neural precursor cells (smNPCs) into midbrain dopaminergic neurons (mDANs) was adapted from an already published method ^55^. Cells were plated onto Geltrex-coated culture dishes, and three distinct media formulations were applied sequentially over specific time intervals. For the initial phase (day 0–8), smNPCs were treated with a medium containing 100 ng/ml FGF8b, 1 µM purmorphamine (PMA), and 200 µM ascorbic acid. From day 8 to day 10, a second medium was introduced, where PMA was reduced to 0.5 µM while maintaining ascorbic acid at 200 µM. Differentiation was completed using a maturation medium applied from day 10 onward, which included 200 µM ascorbic acid, along with 10 ng/ml each of BDNF and GDNF, 500 µM dcAMP, and 1 ng/ml TGFβ3.

### Flow cytometry and FACS

Cells were dissociated using Accumax and incubated at 37 °C for approximately 5–10 minutes. Two volumes of DMEM/F-12 were added and the cell suspension was filtered through a 50 µm strainer into a 15 ml falcon tube. After centrifugation at 300 × g for 3 minutes at room temperature (RT), the pellet was gently resuspended in PBS and transferred to 1.5 ml tubes. Hoechst 33342 (Invitrogen™-H21492) was added to a final concentration of 1 µg/ml, and the samples were rotated at 4 °C for 5 minutes. For dead cell control, one sample was heat-inactivated at 70 °C for 5 minutes. The cells were then washed by centrifugation under the same conditions and resuspended in PBS for further analysis. Flow cytometry was performed using the BD LSRFortessa™ analyzer while for FACS, BD FACSMelody™ Cell Sorter was used for purifying cells transduced with CRISPRa, finally data were analyzed using FlowJo version 10.

### Total RNA extraction, cDNA synthesis and RT-qPCR

Total RNA extraction was performed by first aspirating the medium and adding 600 µl of Lysis Buffer from the Quick-RNA Microprep Kit (Zymo Research) directly to each well. RNA isolation was carried out according to the manufacturer’s guidelines. RNA concentration was determined using a NanoDrop spectrophotometer.

For cDNA synthesis, between 300 ng and 1 µg of RNA, depending on sample availability, was diluted in nuclease-free water to reach a total volume of 27 µl. 13 µl of master mix was then added, containing 1× cDNA synthesis buffer, 0.5 mM dNTPs (Thermo Fisher), 2.5 µM oligo(dT) primer, 40 U/µl Ribolock RNase inhibitor (Thermo Fisher), and 200 U/µl RevertAid Reverse Transcriptase (Thermo Fisher), bringing the final reaction volume to 40 µl. Samples were incubated at 42 °C for 1 hour followed by 70 °C for 10 minutes to terminate the reaction. The cDNA was diluted with nuclease-free water, with dilution factors of 1:10 for 1 µg RNA or 1:3 for 300 ng RNA.

RT-qPCR reactions were set up in a total volume of 20 µl, containing 5 µl of cDNA and 15 µl of master mix prepared with 1×Absolute Blue SYBR Green ROX Low Mix (Thermo Fisher), 500 nM of each primer, and nuclease-free water. Reactions were run in triplicate on MicroAmp™ Fast Optical 96-Well Reaction Plate using an Applied Biosystems 7500 Fast Real-Time PCR System or the QuantStudio 12K Flex Real-Time PCR System under the cycling conditions of 95 °C for 15 minutes, followed by 40 cycles of 95 °C for 15 seconds, 55 °C for 15 seconds, and 72 °C for 30 seconds.

Gene expression analysis was performed using the 2^^−(ΔΔCt)^ method, where ΔΔCt was calculated as: (ΔCt_target − ΔCt_housekeeping) − (ΔCt_target − ΔCt_housekeeping) reference condition. ACTB was used as the reference gene, and expression levels were normalized to the reference condition. Statistical significance was evaluated using either one-sample t-test or ANOVA.

### Immunocytochemistry

Cells were seeded into a PhenoPlate 96-well (PerkinElmer, 6055308). On the day of analysis, cells were fixed with 50 µl of 4% paraformaldehyde for 15 minutes at RT. After fixation, cells were washed three times with PBS containing MgCl₂ and CaCl₂. Permeabilization was performed by incubating the cells for 15 minutes with 75 µl of PBS containing MgCl₂, CaCl₂, 0.4% Triton X-100, 10% goat serum, and 2% BSA. The cells were then washed twice with PBS containing MgCl₂ and CaCl₂, followed by overnight incubation at 4 °C with 50 µl of primary antibody diluted appropriately in PBS with MgCl₂, CaCl₂, 0.1% Triton X-100, 1% goat serum, and 0.2% BSA. After three washes with PBS containing MgCl₂ and CaCl₂, cells were incubated for 1 hour at RT with 50 µl of secondary antibody diluted in the same buffer as the primary antibody. Following three additional washes, cells were incubated for 15 minutes at RT with DAPI (1:1000 dilution) and washed again twice. Fixed and stained cells were stored at 4 °C until analysis.

### High-content imaging

Cells were seeded into a PhenoPlate 96-well (PerkinElmer, 6055308) at 100 000 densities at day 4, while at day 8 of analysis cells were fixed and stained as prior described. The primary antibodies used were rabbit polyclonal anti-ZFHX4 1:100 (Merck-HPA023837), mouse monoclonal anti-Ki67 1:100 (BDPharmingen**-**550609), rabbit polyclonal anti-PH3 1:500 (Merck-06-570), chicken polyclonal anti-MAP2 1:1000 (Abcam-ab5392) with their respective secondary antibodies diluted 1:1000 goat anti-Chicken Alexa Fluor™ 568 (Invitrogen—A11041), donkey anti-Rabbit Alexa Fluor™ 568 (Invitrogen— A10042), goat anti-Chicken Alexa Fluor™ 647(Invitrogen—A21449) and donkey anti-Mouse Alexa Fluor™ 647 (Invitrogen— A31571).

For each well, sixteen image z-stacks were acquired at 20x magnification using the CellVoyager CV8000 High-Content Screening System (Yokogawa). Each z-stack consisted of three focal planes spaced 3.2 µm apart. DAPI fluorescence was excited with a 405 nm laser and collected through a 445/45 nm bandpass filter. Alexa568 was excited using a 561 nm laser and emission was detected through a 600/37 nm bandpass filter. Alexa647 excitation was performed with a 640 nm laser, with emission captured behind a 676/29 nm bandpass filter.

Signal masks for ZFHX4, Ki67, PH3 and MAP2 were created using a pipeline developed in Matlab, by applying thresholding. The signal intensity of each marker, in regions exceeding the threshold, was then calculated and normalized over DAPI.

### Bacterial culture, plasmid extraction and lentivirus production

Glycerol stocks of bacteria carrying the plasmid of interest were cultured from glycerol stocks. Plasmid DNA was then extracted using the NucleoBond Xtra Midi EF kit (Macherey-Nagel) according to the manufacturer’s protocol.

For lentiviral production, 8 × 10⁶ HEK293T cells were seeded into a T75 flask with 15 ml of DMEM (Gibco) supplemented with 10% fetal bovine serum and 1% penicillin-streptomycin. The following day, cells were transfected using a third-generation lentiviral system. A DNA mix consisting of 4 µg pMDG, 2 µg pMDL, 2 µg pREV, and 8 µg of the transfer plasmid was prepared in 200 µl of 1 M CaCl₂ (Sigma), and the volume was adjusted to 800 µl with sterile water. This solution was gently combined with 800 µl of HEPES-buffered saline (Sigma) by adding dropwise while bubbling and incubated at room temperature for 20 minutes. Meanwhile, 16 µl of 25 mM chloroquine was added to the HEK293T cells and incubated for at least 5 minutes. The prepared transfection complex was then added to the cells. After 4–6 hours, the medium was replaced with 14 ml of fresh DMEM. Viral supernatant was collected after 48 hours, centrifuged at 2000 rpm for 10 minutes at 4 °C, and filtered through a 0.45 µm membrane (Sartorius). The resulting viral particles were aliquoted into 1 ml portions and stored at −80 °C.

### CRISPR activation

CRISPR activation (CRISPRa) was used to overexpress ZFHX4. The plasmid pLV hU6-gRNA-hUbC-VP64-dCas9-VP64-T2A-GFP (Addgene #66707) was employed for lentiviral production and constitutive expression of dCas9-VP64 after transduction. The selected gRNAs targeting ZFHX4 and a scramble control were cloned into the backbone of this plasmid using Golden Gate assembly with the BsmBI type IIS restriction enzyme. The cloning of the gRNA sequences was performed by GeneCust. After receiving the new plasmids, One Shot™ Stbl3™ Chemically Competent *E. coli* (Thermo, C7373-03) were transformed following the manufacturer’s instructions. Glycerol stocks were prepared, and plasmid isolation and lentivirus production were performed as previously described.

### Transduction

Lentiviral transduction of differentiating neurons was carried out at specific time points during the protocol, as outlined in Fig. 2A. Cells were pre-seeded in appropriate densities depending on the planned transduction day. On the day of transduction, frozen lentiviral particles were thawed on ice, and fresh differentiation medium was prepared. The medium was removed from each well, and the required volume of virus was added. If the viral volume was less than 1 ml, it was supplemented with medium to reach this amount. Plates were sealed and centrifuged at 300 g for 10 minutes at room temperature to enhance transduction efficiency. Subsequently, additional medium was added to each well to reach a final volume of 2 ml. Media were refreshed in less than 24 hours post-transduction. All lentiviral vectors included a GFP marker to monitor transduction efficiency and viral preparations were titrated beforehand to achieve approximately 80% efficiency, confirmed via GFP expression by flow cytometry.

### TaqMan assay

To evaluate miR-9 expression, the TaqMan™ MicroRNA Assay specific for hsa-miR-9 (ThermoFisher, Assay ID: 000583) was employed together with the U6 snRNA (Assay ID: 001973) as an internal reference. Reverse transcription was performed with the TaqMan™ MicroRNA Reverse Transcription Kit (ThermoFisher), following the supplier’s instructions. Briefly, 3 µl of 5× RT primer was mixed with 5 µl of RNA template containing 1–10 ng of nucleic acid. The mixture was initially incubated at 85°C for 5 minutes and then at 60°C for 5 minutes. Subsequently, 7 µl of a reaction mixture was added, comprising 0.15 µl of 100 mM dNTPs (with dTTP), 1 µl of MultiScribe™ Reverse Transcriptase (50 U/µl), 1.5 µl of 10× RT Buffer, 0.19 µl of RNase inhibitor (20 U/µl), and 4.16 µl of nuclease-free water. The reverse transcription was performed under the following conditions: 30 minutes at 16°C, followed by 30 minutes at 42°C, and a final step at 85°C for 5 minutes.

For RT-PCR, 1.33 µl of the synthesized cDNA was combined with 1 µl of 20× TaqMan™ Small RNA Assay, 10 µl of TaqMan™ Fast Advanced Master Mix, and 7.67 µl of nuclease-free water to a final volume of 20 µl. Amplification was carried out on the Applied Biosystems 7500 Fast Real-Time PCR System. The cycling protocol followed indicated 50°C for 2 minutes, 95°C for 20 seconds, then 40 cycles of 95°C for 3 seconds and 60°C for 30 seconds. Relative quantification was performed using the 2^−ΔΔCt method as previously described.

### CUT&Tag

At day of analysis, cells were detached using Accumax for 5-10 minutes at 37 °C. Two volumes of DMEM/F-12 were added, the cell suspension was filtered through a 50 µm strainer into a 15 ml falcon tube and centrifuged at 300 × g for 3 minutes at RT. The pellet was then resuspended in PBS and transferred to 1.5 ml tubes. DAPI was added to the samples and incubated for 5 minutes on a rotator at 4 °C. After centrifugation with the same conditions, cells were washed twice in PBS containing 2% BSA and were then ready for sorting. For fluorescence-assisted cell sorting (FACS), the BD FACSAria™ III sorter was used. 200 000 cells were obtained for each condition, after sorting. Cells were passed to 1.5 ml tubes and centrifuged for 3 min, 600g at RT, the pellet was then resuspended in PBS and 0.6 ul of formaldehyde 19% were added. Samples were incubated for 2 minutes at RT and 7.5 µl of 1M glycine were added to the cell suspension to stop the cross-linking reaction. CUT&Tag was then performed using the CUT&Tag-IT™ Assay Kit, Anti-Rabbit, 53160, from Active Motif as per manufacturer’s instructions. The antibodies employed were the rabbit polyclonal anti-ZFHX4 (Merck-HPA023837) and the rabbit polyclonal anti-Histone H3 acetyl K27 (ab4729). 1 µg of each antibody was added to 50 µL of solution, resulting in a final concentration of 0.02 µg/ µL.

### Sequencing

RNA quality was assessed with the Agilent RNA 6000 Nano kit on an Agilent 2100 Bioanalyzer. Only samples with a RIN greater than 7 were selected for sequencing. RNA sequencing of ZFHX4 knockdown samples was performed using the Illumina stranded mRNA library preparation kit, generating paired-end reads of 50 base pairs on a NovaSeq6000 platform.

The quality of the CUT&Tag libraries was defined using the Agilent High Sensitivity DNA Kit on the Agilent 2100 Bioanalyzer. The sequencing was carried out in a Nextseq500 machine using paired-end 75 bp read length.

### RNA-seq analysis

For the RNA-seq data from ZFHX4 knock down experiments a snakemake pipeline was used (Köster et al., 2021). This pipeline includes the tools STAR, SAMtools, FastQC, FastQ Screen, AdapterRemoval, Rsubread, DESeq2, ggplot2 and apeglm ^56,57^. For more details about the pipeline, please refer to our repository (RNA-seq folder, RNA-seq_DataAnalysis_TF_KDs.rmd script). The genome version and annotation were GRCh38 release 102. The analysis pipeline was based on the software container https://hub.docker.com/layers/ginolhac/snake-rna-seq/0.4.

### CUT&Tag analysis

The CUT&Tag analysis of H3K27ac and ZFHX4 in mDANs on day 15 was performed using the following tools. Quality control was first conducted for the raw fastq files using FastQC and FastQ. The results were summarized with MultiQC. Sequencing adapters were trimmed using AdapterRemoval and the Reads were then aligned to the reference genome using Bowtie2. The resulting SAM files were converted and sorted using SAMtools, followed by indexing. For peak calling, MACS3 was used, and results were output as bigWig files. Genome version was GRCh38, Ensembl release 104. The analysis was performed using the Docker container <u>ginolhac/snake-cut-and-tag:</u>0.1.

### Primers

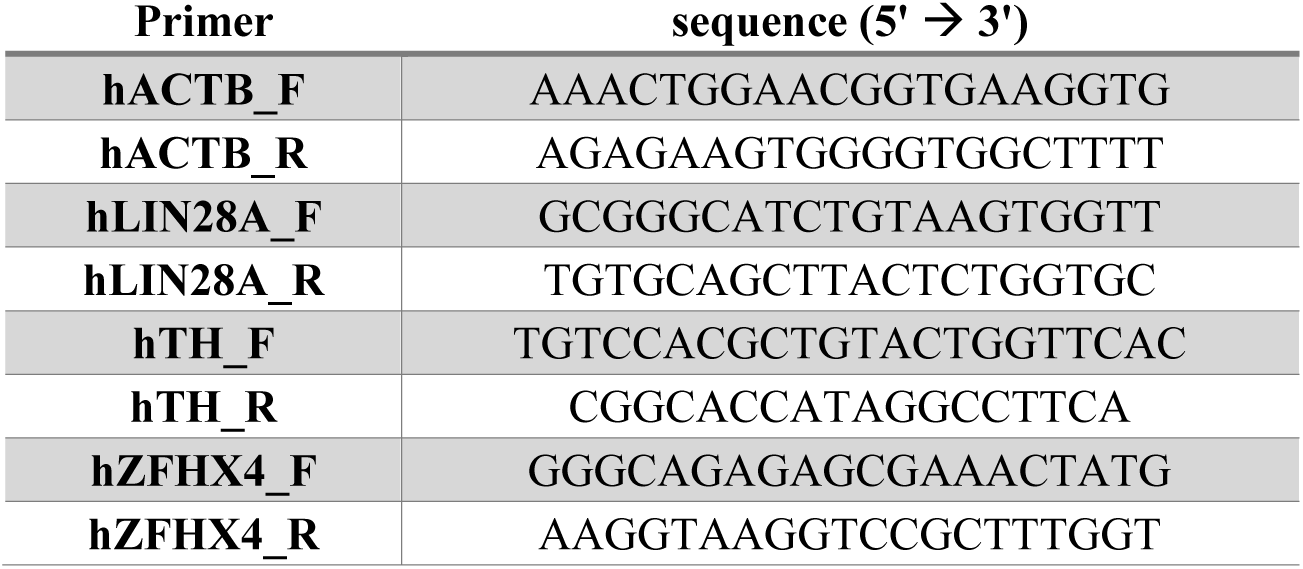

### Primary Antibodies for immunocytochemistry

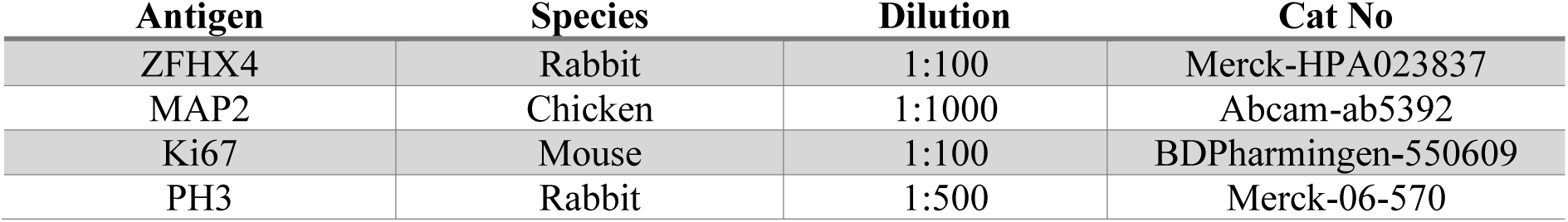

### Bacterial glycerol stocks

All bacterial glycerol stocks are from GeneCopoeia. The different shRNA constructs used were:

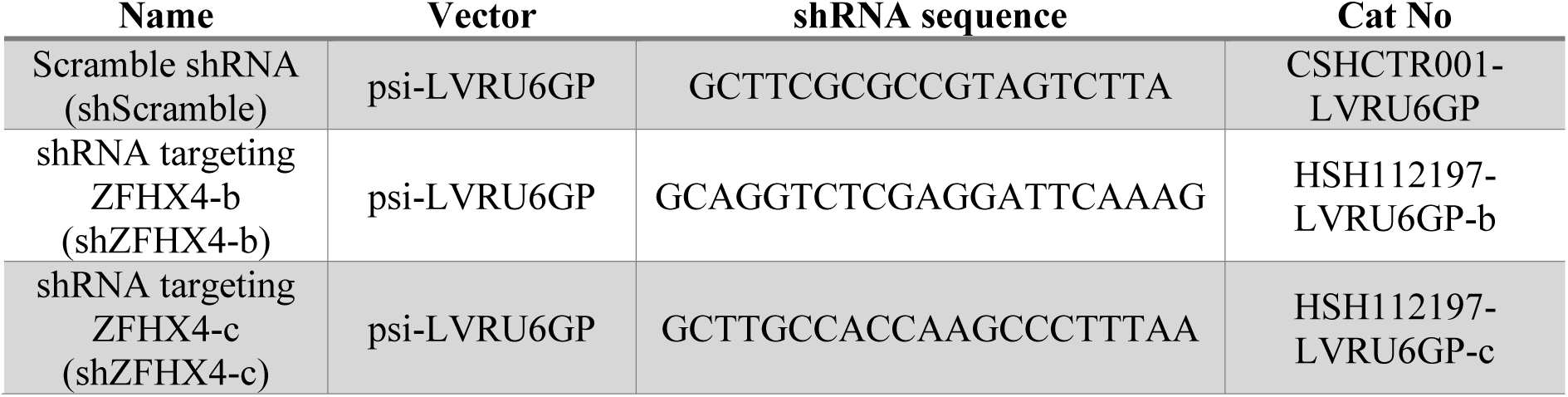

## Data availability

The source RNA-seq and CUT&Tag fastq files have been deposited at https://ega-archive.org/. Additional intermediate files can be provided upon request.

The Snakemake used in this study for the CUT&Tag analysis is available at: https://gitlab.com/uniluxembourg/fstm/dlsm/bioinfo/snakemake-cut-n-tag commit da722539, for the RNA-seq analysis https://gitlab.com/uniluxembourg/fstm/dlsm/bioinfo/snakemake-rna_seq v0.2.3. The code used for RNA-seq data analysis as well as the Matlab code used to analyze the images from the high-content imaging are available at https://github.com/sysbiolux/Valceschini_et_al_2025.

Data used in the preparation of this article were obtained on May, 22, 2023 from the Parkinson’s Progression Markers Initiative (PPMI) database (www.ppmi-info.org/access-data-specimens/download-data), RRID:SCR_006431. For up-to-date information on the study, visit www.ppmi-info.org.

## Acknowledgments

The computational analysis presented in this paper were carried out using the HPC facilities of the University of Luxembourg. We thank the Bioimaging Platform of the Luxembourg Centre for Systems Biomedicine for their support. All schematic representations were created with BioRender.com.

E.V. and L.S. were supported by the AUDACITY grant from the Institute of Advanced Studies at the University of Luxembourg (GENERIC). This work was supported by the Luxembourg National Research Fund within the National Centre of Excellence in Research on Parkinson’s Disease (NCER-PD; FNR/NCER13/BM/11264123) and the PEARL program (FNR/P13/6682797 to R.K.). B.G.R. was funded by the Luxembourg National Research Fund through the PARK-QC doctoral training unit (PRIDE17/12244779/PARK-QC).; L.S., B.G.R., J.O., and D.G. have received funding from Fondation du Pélican de Mie et Pierre Hippert-Faber and Luxembourg Rotary Foundation.

PPMI – a public-private partnership – is funded by the Michael J. Fox Foundation for Parkinson’s Research and funding partners, including 4D Pharma, Abbvie, AcureX, Allergan, Amathus Therapeutics, Aligning Science Across Parkinson’s, AskBio, Avid Radiopharmaceuticals, BIAL, Biogen, Biohaven, BioLegend, BlueRock Therapeutics, Bristol-Myers Squibb, Calico Labs, Celgene, Cerevel Therapeutics, Coave Therapeutics, DaCapo Brainscience, Denali, Edmond J. Safra Foundation, Eli Lilly, Gain Therapeutics, GE HealthCare, Genentech, GSK, Golub Capital, Handl Therapeutics, Insitro, Janssen Neuroscience, Lundbeck, Merck, Meso Scale Discovery, Mission Therapeutics, Neurocrine Biosciences, Pfizer, Piramal, Prevail Therapeutics, Roche, Sanofi, Servier, Sun Pharma Advanced Research Company, Takeda, Teva, UCB, Vanqua Bio, Verily, Voyager Therapeutics, the Weston Family Foundation and Yumanity Therapeutics.

The transcriptomic and epigenomic data supporting the conclusions of this article are available in the IHEC EpiATLAS resource repository, https://epigenomesportal.ca/ihec/, (International Human Epigenome Consortium, EpiATLAS - a reference for human epigenomic research, in preparation)

## Supplementary Figure Legends

**Supplementary Figure 1:**
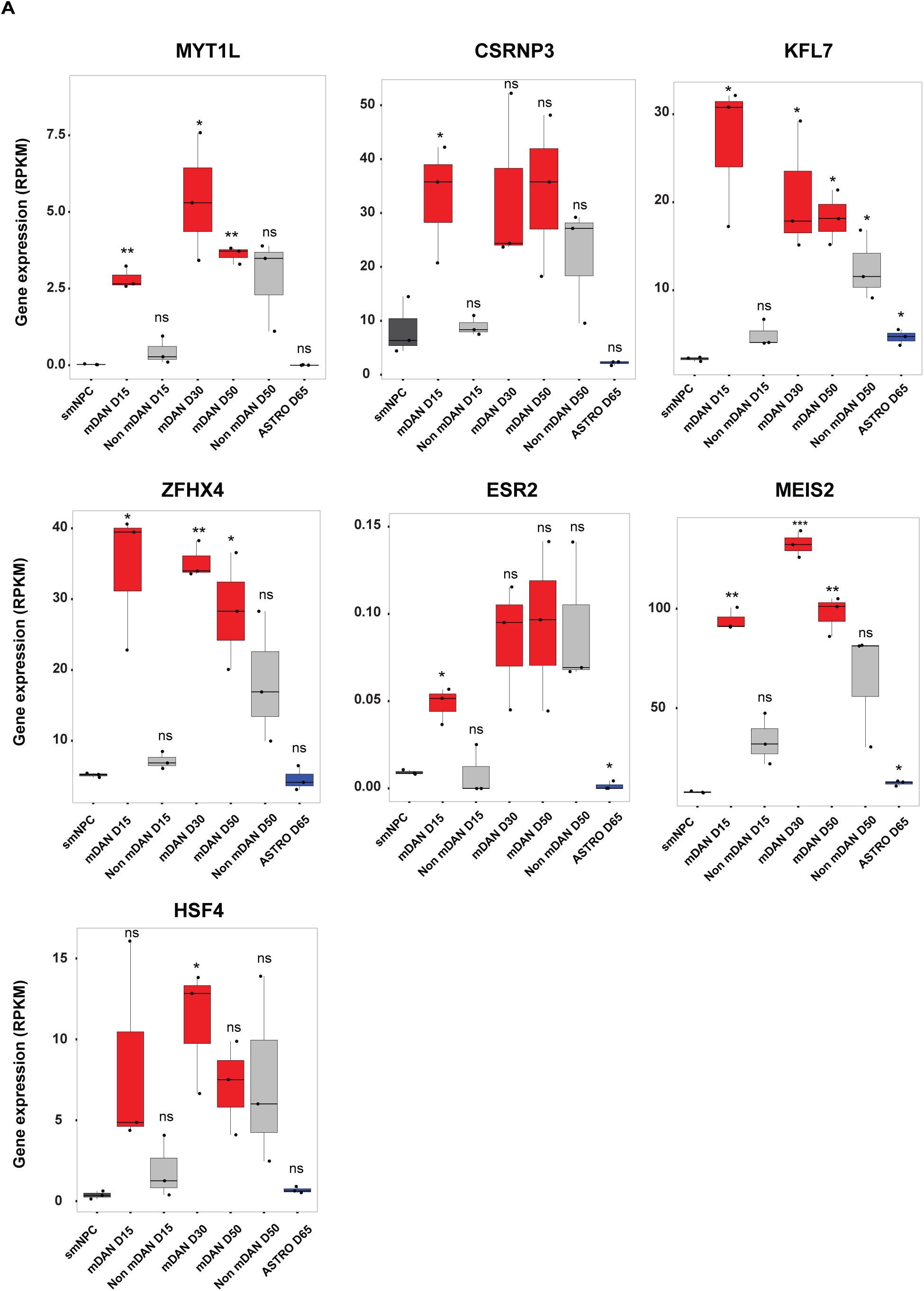
Expression of 7 TFs across mDAN differentiation. A) Expression dynamics of the 7 candidate TFs during mDAN differentiation of the TH-Rep1 cell line.

**Supplementary Figure 2:**
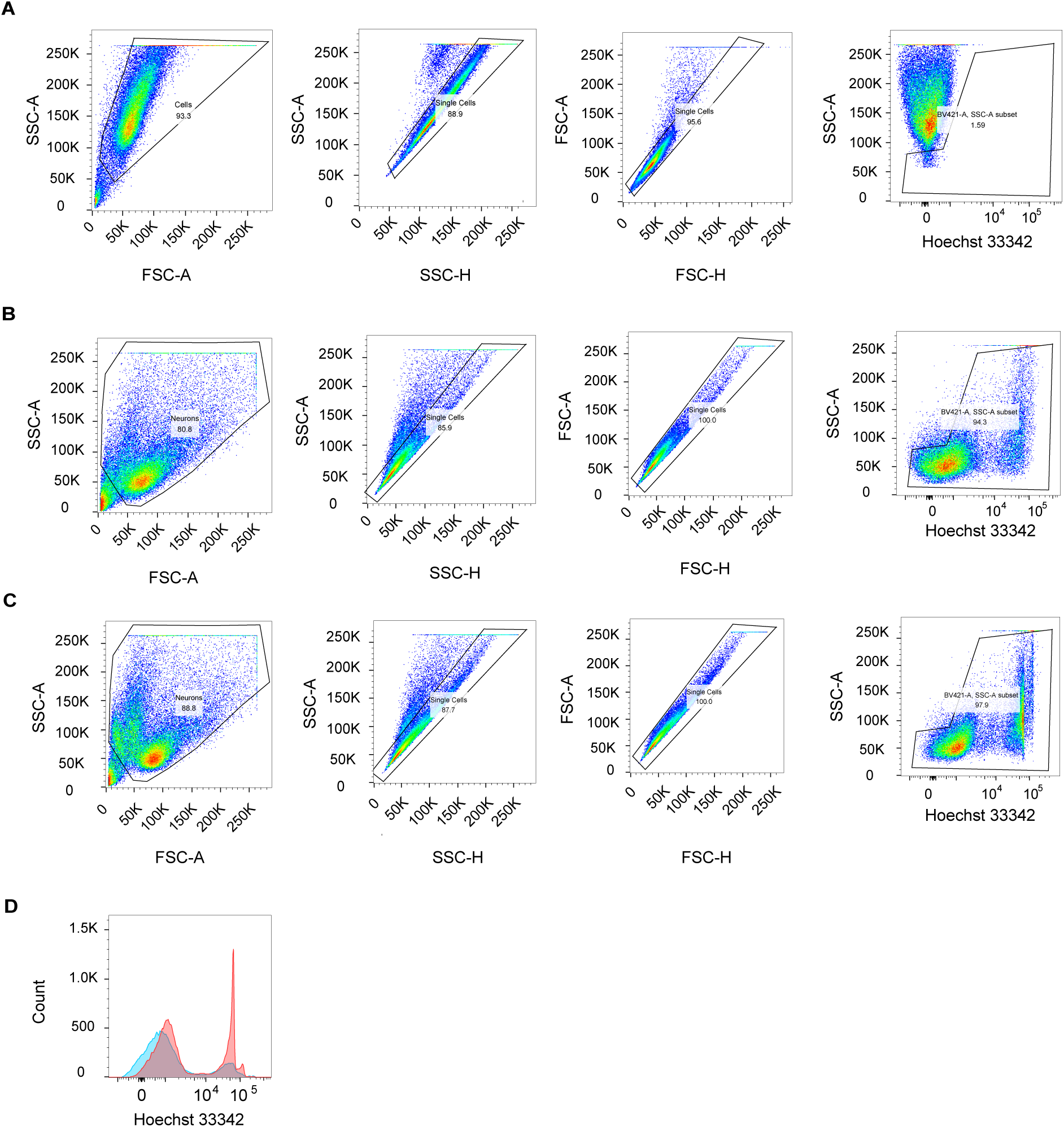
Gating strategy for the cell cycle analysis upon ZFHX4 KD. A) Representative gating of a dead mDAN control culture used to define the non-viable population. B) Gating strategy for the shSCRAMBLE condition. C) Gating strategy for the shZFHX4 condition. All samples were stained under the same conditions.

